# A novel transposable element based authentication protocol for *Drosophila* cell lines

**DOI:** 10.1101/2021.08.16.456580

**Authors:** Daniel Mariyappa, Douglas B. Rusch, Shunhua Han, Arthur Luhur, Danielle Overton, David F. B. Miller, Casey M. Bergman, Andrew C. Zelhof

## Abstract

*Drosophila* cell lines are used by researchers to investigate various cell biological phenomena. It is crucial to exercise good cell culture practice. Poor handling can lead to both inter- and intraspecies cross-contamination. Prolonged culturing can lead to introduction of large- and small-scale genomic changes. These factors, therefore, make it imperative that methods to authenticate *Drosophila* cell lines are developed to ensure reproducibility. Mammalian cell line authentication is reliant on short tandem repeat (STR) profiling, however the relatively low STR mutation rate in *D. melanogaster* at the individual level is likely to preclude the value of this technique. In contrast, transposable elements (TE) are highly polymorphic among individual flies and abundant in *Drosophila* cell lines. Therefore, we investigated the utility of TE insertions as markers to discriminate *Drosophila* cell lines derived from the same or different donor genotypes, divergent sub-lines of the same cell line, and from other insect cell lines. We developed a PCR-based next-generation sequencing protocol to cluster cell lines based on the genome-wide distribution of a limited number of diagnostic TE families. We determined the distribution of five TE families in S2R+, S2-DRSC, S2-DGRC, Kc167, ML-DmBG3-c2, mbn2, CME W1 Cl.8+, and OSS *Drosophila* cell lines. Two independent downstream analyses of the NGS data yielded similar clustering of these cell lines. Double-blind testing of the protocol reliably identified various *Drosophila* cell lines. In addition, our data indicate minimal changes with respect to the genome-wide distribution of these five TE families when cells are passaged for at least 50 times. The protocol developed can accurately identify and distinguish the numerous *Drosophila* cell lines available to the research community, thereby aiding reproducible *Drosophila* cell culture research.

## Introduction

As of 2018, the estimated of the number of publications using all cell culture studies is ∼2 million (Bairoch 2018). However, problems with reproducibility and authenticity hamper their use (Almeida *et al*. 2016). Poor culture practices in individual laboratories has led to many cases of inter- and intraspecies cross-contamination (Capes-Davis *et al*. 2010). Additionally, prolonged passaging can lead to large- and small-scale genomic changes due to *in vitro* evolution that cause sub-lines of the same cell line to vary among laboratories (Ben-David *et al*. 2018; Liu *et al*. 2019). For example, extensive passaging (>50 passages) of viral-transformed human lymphoblastoid cell lines is associated with increased genotypic instability (Oh *et al*. 2013). Likewise, long term passaging of mammalian cell lines is known to lead to increased single nucleotide variations (Pavlova *et al*. 2015), reduced differentiation potential (Yang *et al*. 2018) and changes in the karyotype (Wenger *et al*. 2004). To overcome these inconsistencies in experiments across laboratories when using human cell lines, the American National Standards Institute and the American Type Culture Collection (ANSI/ATCC ASN-002) have provided a standard for vertebrate cell culture work. Moreover, the NIH offers guidelines for authenticating key research resources that have been endorsed by several major journals (ATCC 2011; NIH 2015).

Though most of above-mentioned problems and solutions relate to mammalian cell culture practice, a significant number of laboratories use *Drosophila* cells for basic research. *Drosophila* cell lines are used by researchers to investigate a myriad of cellular processes including receptor-ligand interactions (Ozkan *et al*. 2013), cellular signaling (Albert and Bokel 2017), circadian biology (Albert and Bokel 2017), metal homeostasis (Mohr *et al*. 2018), cellular stress response (Aguilera-Gomez *et al*. 2017), neurobiology (Tsuyama *et al*. 2017), innate immunity (Nonaka *et al*. 2017), and functional genomics (Albert and Bokel 2017), as well as being used extensively for gene editing by CRISPR Cas9 technology (Luhur *et al*. 2018). Furthermore, as part of the modENCODE project, the transcriptional and chromatin profiles of a large panel of *Drosophila* cell lines were determined to facilitate studies on gene function and expression (Cherbas *et al*. 2011; Kharchenko *et al*. 2011). However, currently there are no protocols available to authenticate *Drosophila* cell lines. In addition, the effects of long-term passaging on *Drosophila* cell lines have not been formally investigated despite evidence for extensive changes from wild-type ploidy and copy number in many *Drosophila* cell lines (Zhang *et al*. 2010; Lee *et al*. 2014), implying that insect cells can potentially exhibit genomic changes in culture like their mammalian counterparts.

Human cell line authentication guidelines recommend short tandem repeat (STR) profiling as the method of choice for routine cell typing, although approaches using genomic techniques yield more comprehensive information (Almeida *et al*. 2016). The use of STR profiling as the preferred method to authenticate human cell lines is based on high STR allelic diversity among the donors for different cell lines, relatively low cost, stability of using STR markers, and the historical availability of methods to assay STR variants during the development of human cell line authentication protocols. There are a number of limitations with the STR approach. The ANSI/ATCC ASN-002 standard for typing human cell lines with STRs is over 100 pages long and requires careful implementation for proper interpretation. Moreover, STR-based methods for human cell line authentication are primarily designed to discriminate cell lines derived from different donors, but are less powerful for discriminating cell lines or sub-lines from the same donor genotype.

Development of cell line authentication protocols requires understanding the genome biology of a species, the specific characteristics of the most widely used cell lines in that research community, and how these features can be used to leverage cost-effective modern genomic technologies. In *Drosophila*, the majority of widely-used cell lines have been derived from a limited number of donor genotypes. Coupled with the low STR mutation rate in *Drosophila* relative to humans (Schug *et al*. 1997), the use of STR profiling for discriminating different *Drosophila* cell lines is likely to be limited. In contrast, it is well-established that transposable elements (TE) are highly polymorphic among individual flies (Charlesworth and Langley 1989) and that *Drosophila* cell lines have an increased TE abundance relative to whole flies (Potter *et al*. 1979). These properties, together with the large number of potential insertion sites across the genome and stability of TE insertions at individual loci, suggest that TE insertions should theoretically be useful markers to simultaneously discriminate *Drosophila* cell lines made from different donor genotypes as well as from the same donor genotype, including divergent sub-lines of the same cell line. Han et. al (2021) recently tested this prediction and demonstrated that genome-wide TE insertion profiles can reliably cluster different *Drosophila* cell lines from the same donor genotypes and discriminate cell lines from different donor genotypes, while also preserving information about the laboratory of origin. A minimal subset of six active TE families (*297, copia, mdg3, mdg1, roo* and *1731*) was also determined to have essentially the same discriminative power as the genome-wide dataset (Han *et al*. 2021).

Based upon these findings, we investigated if the genome-wide distribution of these six TE families could form the basis for a reliable protocol to authenticate *Drosophila* cell lines. As noted earlier, several of the modENCODE cell lines are extensively used to study genomic and cell biological processes (Cherbas *et al*. 2011; Kharchenko *et al*. 2011). These cell lines are also amongst the most widely-ordered cell lines from *Drosophila* Genomics Resource Center (DGRC). Therefore, we used six modENCODE lines derived from various *D. melanogaster* developmental stages: S2R+, S2-DRSC, Kc167 (embryonic origin); ML-DmBG3-c2 (L3 larval CNS origin); mbn2 (larval circulatory system origin); and CME W1 Cl.8+ (wing disc origin) in our analysis. Two other non-modENCODE cell lines – S2-DGRC and OSS (ovarian somatic sheath) – that are ordered frequently from the DGRC were also included.

Here we present data supporting the utility of a genomic TE distribution (gTED) protocol to authenticate *D. melanogaster* cell lines. The developed gTED protocol was able to generate distinct TE genomic distribution signatures for all the cell lines tested. Moreover, using the gTED protocol we were able to authenticate blinded samples from the *Drosophila* research community, thus validating the protocol. Moreover, the gTED signatures of up to 50 passages of S2R+ cells do not cluster in a passage-dependent manner, indicating that this protocol could be used to authenticate cell lines with up to 50 passages. Moving forward, we aim to expand the repertoire of cell lines assessed for their TE genomic distribution. We now have a protocol that can be adopted by the *Drosophila* research community to authenticate their cell lines and provide the necessary standards as per NIH guidelines.

## Materials and Methods

### Drosophila cell lines and genomic DNA extraction

Our protocol development included six modENCODE lines derived from various *Drosophila* developmental stages: embryonic - S2R+ (DGRC #150, CVCL_Z831), S2-DRSC (DGRC #181, CVCL_Z992), Kc167 (DGRC #1, CVCL_Z834); L3 larval CNS origin - ML-DmBG3-c2 (DGRC #68, CVCL_Z728); larval circulatory system origin - mbn2 (DGRC #147, CVCL_Z706); and wing disc origin - CME W1 Cl.8+ (DGRC #151, CVCL_Z790) (Table 1). Two other non-modENCODE cell lines – S2-DGRC (DGRC #6, CVCL_TZ72) and OSS (ovarian somatic sheath, DGRC #190, CVCL_1B46), were also included in the protocol development phase. The S2R+, S2-DRSC, S2-DGRC, mbn2 cells were cultured in the Shields and Sang M3 medium (Sigma, Cat#: S8398) supplemented with 10% fetal bovine serum (FBS, Hyclone, GE Healthcare), bactopeptone (Sigma) and yeast extract (Sigma) M3+BPYE+10%FBS. ML-DmBG3-c2 cells were cultured in M3 + BPYE + 10% FBS with 10 µg/ml insulin (Sigma-Aldrich) while CME W1 Cl.8+ cells required M3 + 2% FBS + 5 µg/ml insulin + 2.5% fly extract containing medium. OSS cells were cultured in M3 + 10% FBS + 10% fly extract with 60 mg L-glutathione (Sigma-Aldrich, Cat#: G6013) and 10 µg/ml insulin (Sigma-Aldrich, Cat#: I9278). Kc167 cells were cultured in CCM3 medium (Hyclone, Cat#: SH30061.03). To extract total genomic DNA, cells were cultured to confluency, harvested by pipetting, centrifuged and washed once with phosphate-buffered saline (PBS). Genomic DNA (gDNA) was extracted from the PBS washed pellet using the Zymo Quick-DNA™ MiniprepPlusKit (Cat#: D4068/4069), using 1 column for every 10 million cells. Genomic DNA was generated for triplicate samples of all cell lines in order to investigate the reproducibility of our protocol as well as to detect and mitigate potential mislabeling of individual samples during the project.

**Table 1:** Summary of transposable element (TE) insertions detected by gTED. The total number TE insertions that were detected in each of the listed cell lines is presented as a mean (n=3) of the samples analyzed. SD=standard deviation, CNS: Central Nervous System

| ***Cell line*** | ***Tissue source*** | ***DGRC Stock Number*** | ***Cellosaurus ID*** | ***Number of TE insertions***<br>***Mean (+/- SD)*** |
| --- | --- | --- | --- | --- |
| S2R+ | Embryo | 150 | CVCL_Z831 | 1009 (± 30.4) |
| S2 DGRC | Embryo | 6 | CVCL_TZ72 | 704 (± 3.2) |
| mbn2 | Larval circulatory system | 147 | CVCL_Z706 | 633 (± 6.4) |
| S2 DRSC | Embryo | 181 | CVCL_Z992 | 530 (± 14.8) |
| Kc167 | Embryo | 1 | CVCL_Z834 | 516 (± 18.3) |
| OSS | Adult ovary | 190 | CVCL_1B46 | 404 (± 8.5) |
| CME-W1-CI.8+ | Larval wing disc | 151 | CVCL_Z790 | 309 (± 11.1) |
| ML-DmBG3-c2 | Larval CNS | 68 | CVCL_Z728 | 227 (± 4.7) |

### Blinded samples

External blinded samples from eight cell lines were obtained as triplicates of frozen genomic DNA samples extracted from insect cell lines from Dr. Sharon Gorski, British Columbia Cancer Research Centre, Vancouver, Canada and the *Drosophila* RNAi Screening Center, Harvard University (Table 2). The identities of the external samples sent to DGRC were blinded by the sample donors. For internal blinded samples, genomic DNA was extracted from three cell lines in triplicate (Table 2). The identities of the internal samples were blinded from the team members involved in library preparation and downstream analyses. Genomic DNA for both the external and internal blinded samples was extracted as per the protocol described above. The team members involved in library preparation and downstream analyses were blind to the identity and replicates of each sample.

**Table 2:** List of blinded samples processed. Blinded samples were donated by external (*Drosophila* RNAi Screening Center and Dr. S. Gorski) or generated internally. The identifications were made upon processing the sample through the genomic TE distribution pipeline followed by computational analysis. No ID: The genomic TE signatures of the cell lines did not match with any of the lines analyzed to provide a positive identification. *A. a*: cell line derived from *Aedes aegypti*; *A. g*: cell line derived from *Anopheles gambiae*.

| *Sample label* | *Source* | *Identification with gTED pipeline* | *Confirmation* |
| --- | --- | --- | --- |
| DRSC_Blinded_1-3 | DRSC | Kc167 | Kc167 |
| DRSC_Blinded_4-6 | DRSC | No ID | *A. g* |
| DRSC_Blinded_7-9 | DRSC | No ID | *A. a* |
| DRSC_Blinded_10-12 | DRSC | Kc167 | Kc167 |
| DRSC_Blinded_13-15 | DRSC | S2R+ | S2R+ |
| DRSC_Blinded_16-18 | DRSC | S2 | S2 |
| SGLab_Blinded_1-3 | Gorski Lab | mbn2 | mbn2 |
| SGLab_Blinded_4-6 | Gorski Lab | S2 | S2 |
| DGRC_Blinded_A | Internal | No ID | 1182-4H |
| DGRC_Blinded_B | Internal | No ID | Ras[V12];wts [RNAi] |
| DGRC_Blinded_C | Internal | No ID | delta I(3)mbt-OSC |

### Passage experiment

S2R+ cells were plated at 1 × 10^6^ cells per ml at every passage. A single passage experiment was performed wherein cells were passaged every 2-3 days and replicates of the passages were frozen at the 1^st^, 10^th^, 20^th^, 30^th^, 40^th^ and 50^th^ passages with the cell concentrations between 2.5 – 8.6 × 10^6^ cells per ml. Triplicate genomic DNA samples from each passage was extracted as described above.

### Primer design

Six TE families shown by Han et. al (2021) to be sufficient to identify *Drosophila* cell lines based on WGS data were used as initial candidates for primer design. These six TE families are all long terminal repeat (LTR) retrotransposons, which insert as full-length elements containing an identical LTR that provides a reliably known junction for PCR at each terminus of the TE (Smukowski Heil *et al*. 2021). Primer design was based upon the protocol outlined in Figure 1, involving a two-step PCR (Reaction A/B and Reaction A/B Nest PCR). Each step required one primer to be within the TE at either end (one for Reaction A at the 5’ of the TE and one for Reaction B at the 3’ of the TE). Additionally, primers for Reaction A/B and Reaction A/B Nest PCR needed to have low similarity. Based on these requirements, the general workflow for designing PCR primers for six diagnostic TE families for the eight focal cell lines was as follows:

1. *Generate consensus sequences for LTRs of candidate TE families*.
  a. Whole genome sequencing (WGS) data from (Zhang *et al*. 2010; Lee *et al*. 2014) and (Han *et al*. 2021) for all focal cell lines were mapped against TE canonical sequences and merged into a single BAM file.
  b. Variants were called on the merged BAM file and a VCF file was generated using bcftools call (v1.9).
  c. Full length consensus sequences for all six TE families from VCF file was generated using bcftools consensus (v1.9) with variable sites encoded as ambiguities.
  d. Both the 5’ and 3’ LTRs from the full-length TE consensus sequence for each family were extracted.
2. *Detect the first round of primer candidates*. Primers for nested PCR were detected with primer3 (v2.5.0) (https://github.com/primer3-org/primer3) using the following parameters: PRIMER_LIBERAL_BASE=1; PRIMER_MAX_NS_ACCEPTED=1; PRIMER_NUM_RETURN=10; PRIMER_GC_CLAMP=1; PRIMER_DNA_CONC=25; PRIMER_SALT_MONOVALENT=50; PRIMER_MIN_TM=60; PRIMER_OPT_TM=62; PRIMER_MAX_TM=65; PRIMER_SALT_DIVALENT=2; PRIMER_DNTP_CONC=0; PRIMER_TM_FORMULA=1 PRIMER_OPT_SIZE=22; PRIMER_MIN_SIZE=18; PRIMER_MAX_SIZE=25; PRIMER_MIN_GC=40; PRIMER_MAX_GC=60; PRIMER_PRODUCT_SIZE_RANGE=75-100 150-250 100-300 301-400 401-500 501-600 601-700 701-850 851-1000.
3. *Detect the second round of non-overlapping primer candidates* The same parameters as in the previous round of primer design were used, with the additional specification that the primers designed in the first round were added to a “mispriming library” to exclude these regions for primer prediction in the second round of primer candidates.
4. *Finalize primers from both rounds of primer candidates* The final primers for Reaction A/B PCR and Reaction A/B Nest PCR were selected from the candidate list from both rounds of primer design. Specifically, one primer was selected for Reaction A/B PCR from either round of primer design, then another primer was selected for Reaction A/B Nest PCR from the other round of primer design.

**Figure 1:**
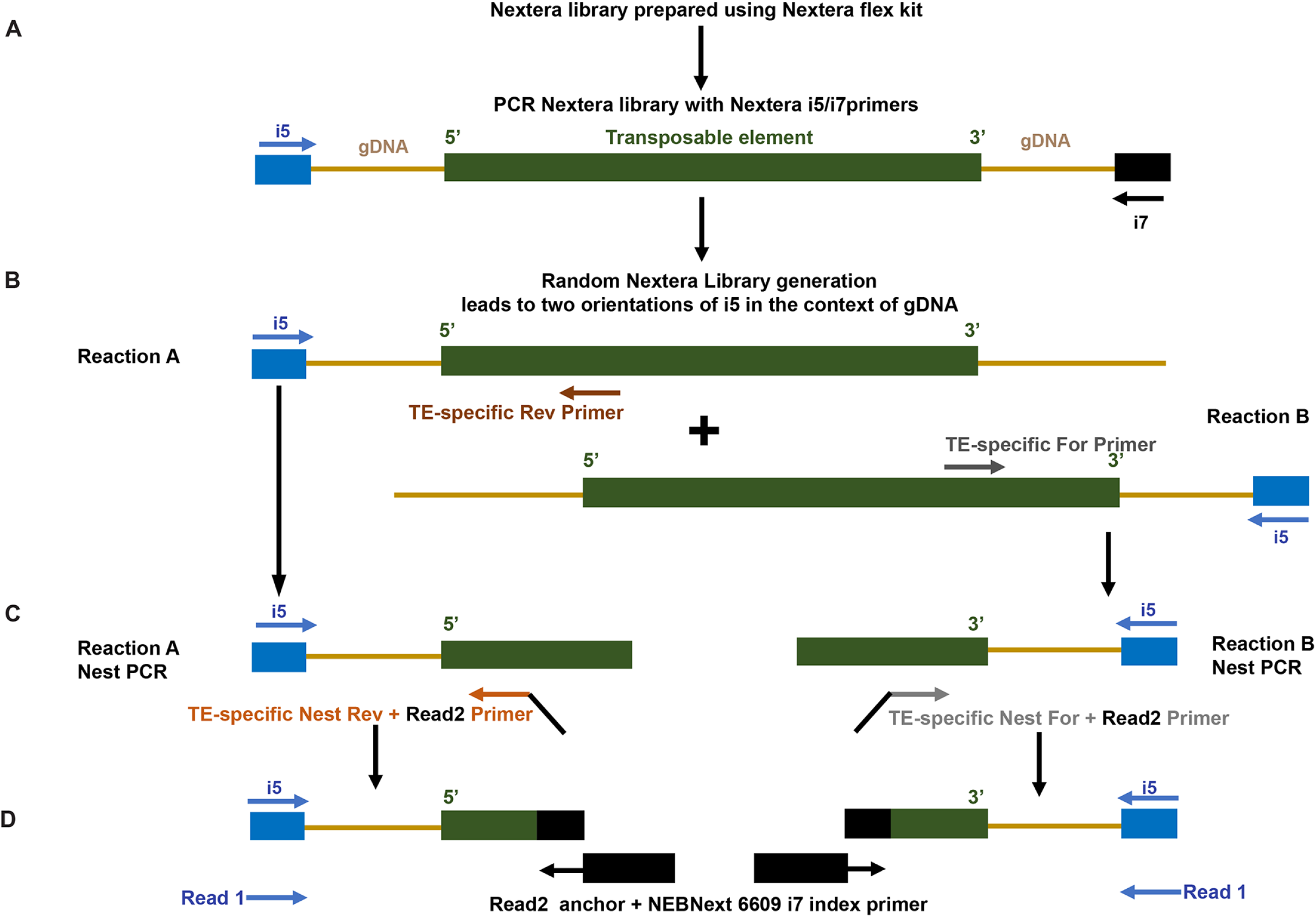
Protocol used for generating libraries to establish genomic transposable element distribution signatures. **A**) Fragmented genomic DNA (gDNA; light brown lines) from the Nextera libraries containing TEs (green bar) and flanking gDNA were amplified with the randomly oriented i5 (blue arrow) and i7 (black arrow) primers. **B**) Reactions A and B involved amplification with the i5 primer oriented in either direction with respect to the TE, in combination either with TE-specific Reverse (dark brown arrow) and Forward (dark grey arrow) primers, respectively. **C**) The Nest PCR reactions amplified from within the products of the respective Reactions A and B using the i5 primer and either the TE-specific Nest Reverse (light brown arrow) or TE-specific Nest Forward (light grey arrow) primers. Read 2 anchors were added onto both the Nest PCR primers. **D**) The final amplification step was performed with the i5 primer and the Read 2 anchor with the i7 index primer (black box). The reads from the genome sequences flanking the TE are designated as Read 1; the reads internal to the TE are designated Read 2.

Final adjustments to the primer locations were made based on testing the respective primer pairs. The full list of primers used in the study are listed in Table S1.

#### Nextera library preparation and nested PCR protocol

Nextera libraries were constructed for all the genomic DNA samples by using Nextera DNA Flex Library Prep Kit (Illumina, Cat#: 20018705) (Figure 1A). Then, the Nextera libraries were diluted into 1nM, and 5 μl of each was used as the template for the TE library construction. To amplify the fragments with the TE-specific genomic context, two separate multiplex PCRs were performed (Reactions A and B, Figure 1B) using TE-specific primers for all six families simultaneously in combination with the Illumina i5 primer. For Reactions A and B, two sets of primers (Forward and Reverse) were designed within the two LTRs of each of the TEs as detailed above. Since the generation of the Nextera library is not direction specific, DNA fragments can orient in either direction with respect to the i5 adaptor thus allowing for detection at either ends of the TE by amplification with the Illumina i5 primer with a TE-specific primer. Therefore, this PCR step amplified the DNA fragments containing the 5’ (Reaction A, Reverse primer) or 3’ (Reaction B, Forward primer) flanking regions of the TEs. A second nested PCR was performed to enrich for the TE-genomic DNA junctions, utilizing nested primers from within the Reactions A and B with the i5 adaptor (Figure 1C). Both Nest PCR primers contained a specific overhang region (5’ GTTCAGACGTGTGCTCTTCCGATCT 3’) to facilitate addition of the index in the next PCR step. The final step was the Index PCR, which was performed to add the i7 adaptor and index by using the kit NEBNext® Multiplex Oligos for Illumina (cat: 6609S). Briefly, equal volumes of the products of Reactions A and B Nest PCRs containing either the TE 5’ and 3’ flanking regions were combined and used as the template. The Index PCR was performed by using the Illumina i5 primer and the NEBNext® Multiplex Oligos to add i7 adaptor and index (Figure 1D). Finally, the TE libraries were constructed with both i5 adaptors (added by Nextera library construction), i7 adaptors and indexes (added by the Index adding PCR).

## Protocol

Step 1:

- Nextera libraries are made by following standard protocol.
- Each library is diluted to 1nM.

Step 2: Reaction A/B (Two sets of reactions)

Reaction A: Primers: i5 + TE Reaction A Rev (To amplify the 5’ flanking region of TE gene)

Reaction B: Primers: i5 + TE Reaction B For (To amplify the 3’ flanking region of TE gene)

### 2.1 PCR reagents

|  |  |
| --- | --- |
| 5X Phusion buffer | 10 µl |
| 100 mM dNTP mix | 0.5 µl |
| 100 uM i5 Primer | 0.5 µl |
| 100 uM Reaction A/B (Rev/For) | 0.5 µl |
| Phusion polymerase | 0.5 µl |
| 1nM Library | 5.0 µl |
| ddH<sub>2</sub>O | 33 µl |
| Total | 50 µl |

### 2.2 PCR settings

2.2 PCR settings:
|  |  |  |
| --- | --- | --- |
| 98°C | 30 sec |  |
| 98°C | 10 sec |  |
| 65°C | 30 sec | 10 cycles |
| 72°C | 60 sec |  |
| 72°C | 5 min |  |
| 4°C | Hold |  |

### 2.3 Cleaned with 0.9X AMPure XP beads, washed with 80% ethanol twice, and elute with 40 μl Elution Buffer (EB)

**Step 3:** Nest PCR (Two sets of reactions)

Set 1: Primers: i5 + TE Reaction A Nest PCR Reverse (Template: Reaction A products)

Set 2: Primers: i5 + TE Reaction B Nest PCR Forward (Template: Reaction B products)

### 3.1 PCR reagents

|  |  |
| --- | --- |
| 5X Phusion buffer | 10 µl |
| 100 mM dNTP mix | 0.5 µl |
| 100 uM i5 Primer | 0.5 µl |
| 100 uM NestPCR Reaction A/B (Rev/For) | 0.5 µl |
| Phusion polymerase | 0.5 µl |
| Reaction A/B products | 38 µl |
| Total | 50 µl |

### 3.2 PCR setting

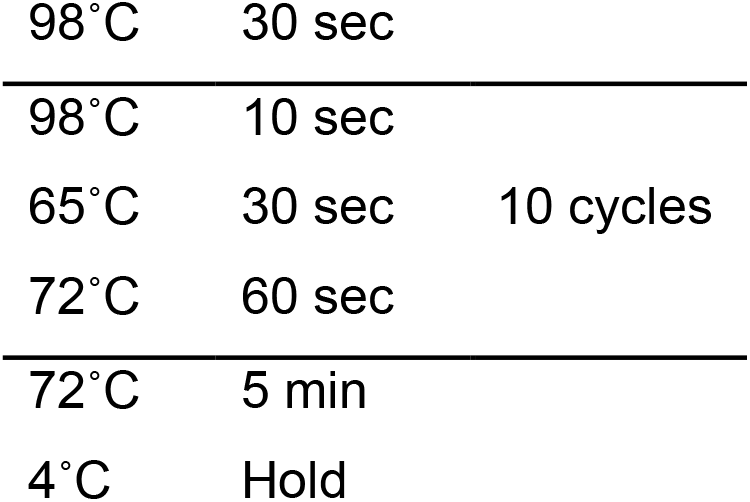

### 3.3 Cleaned with 0.9X AMPure XP beads, wash with 80% ethanol twice, and eluted with 19 μl EB

**Step 4:** Index adding PCR with NEBNext 6609 Primers

### 4.1 PCR reagents

|  |  |
| --- | --- |
| 5X Phusion buffer | 10 µl |
| 100 mM dNTP mix | 0.5 µl |
| 100 uM i5 Primer | 0.5 µl |
| NEBNext 6609S Primer | 5 µl |
| Phusion polymerase | 0.5 µl |
| Nest PCR Reaction A + B products | 33.5 µl |
| Total | 50 µl |

### 4.2 PCR settings

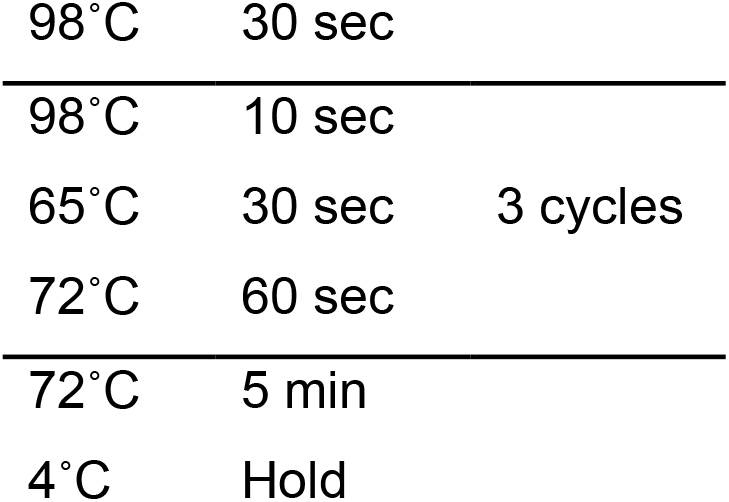

### 4.3 Cleaned with 0.8X AMPure XP beads, washed with 80% ethanol twice, and eluted with 32 μl of EB

#### Sequencing

Paired end sequencing was performed on an Illumina NextSeq 500 with a 150-cycle midi-cycle kits. The first read in a pair (Read 1, R1) corresponds to flanking genomic DNA; the second read in a pair (Read 2, R2) corresponds to TE sequence. Raw sequencing data was submitted to SRA (SRP323476).

#### Sample Processing and Transposable Element Identification

Reads were trimmed for adapters and low quality using Trimmomatic (v0.38; ILLUMINACLIP:adapters.fa:3:20:6 LEADING:3 TRAILING:3 SLIDINGWINDOW:4:20 MINLEN:40). By design, R2 reads occur inside the TE and can be used to demultiplex individual fragments by TE of origin from a multiplex PCR. To do this, R2 reads were aligned to a database of the consensus sequences used for primer design of the relevant TEs using Bowtie2 (v2.3.5.1); the corresponding R1 reads from the same fragment were then demultiplexed into TE specific bins based on the best alignment of R2. R1 reads were then mapped with Bowtie2 (--local -k 2) to the complement and reverse-complement *D. melanogaster* genome (version 6.30) in which the TEs were N-masked (Figure 2; red plus green reads). Masking was performed by searching consensus transposable elements sequences against the *D. melanogaster* genome (version 6.30) using NCBI blastn (version 2.2.26) with the following parameters: -a 10 -e 1e-100 -F “m L” -U T -K 20000 -b 20000 -m 8. R1 reads that did not map with a uniquely best match to the genome were subsequently excluded. Simultaneously, the R1 reads were mapped to the TE consensus sequences. The initial goal was to identify any valid junction where we could explicitly identify the transition from a unique genomic context into a TE, aka a TE junction (Figure 2; green reads). For a R1 read to identify a junction, the local alignment to the genome and the TE must be congruent such that the entire read was accounted for (+/-2 bases). Valid junctions were defined such that multiple independent reads with independent start sites in the genome all identify the same breakpoint. To improve the sensitivity, all the data from all the different samples was combined for junction identification. A valid junction had to have at least 12 reads with 4 distinct start positions. Once the junctions were identified, 300 bp of genomic sequence outside and juxtaposed to the TE junction were isolated, which would include either 5’ or 3’ or both ends of the inserted TE (Figure 2).

**Figure 2:**
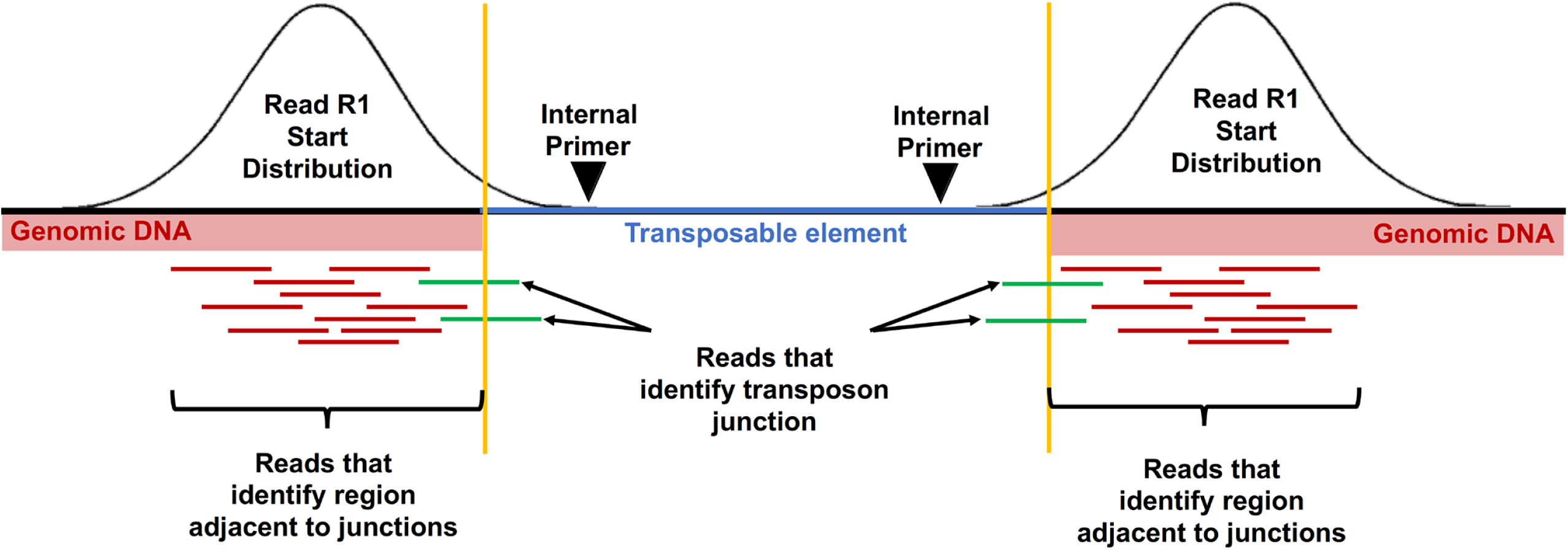
Read mapping strategy used to generate genomic transposable element distribution signatures. Read 1 (R1) reads from demultiplexed fragments were used to identify the transposon junctions (green) from the set of all R1 reads. The schematic represents R1 reads at junctions on either end (5’ or 3’) of a TE. The number of reads that specifically identify a junction is relatively small compared to the total number of reads near the junction. Variation in sequencing depth and subtle differences in the insert sizes produced by the Nextera library could cause junctions to be missed if only explicit junction calls are used. To avoid these issues, after the junctions have been identified, a 300 bp region of genomic sequence flanking the transposon is used to quantify the number of R1 reads (red) associated with that junction.

#### Clustering and Visualization

Read datasets were analyzed in their entirety or by random sub-sampling using vsearch (v2.14.2) (Rognes *et al*. 2016) down to 10 million reads, in order to control for sequencing depth and explore how many reads were necessary per cell line to produce reliable results. Read counts from sub-sampled datasets mapped to dm6 in the 300 bp intervals adjacent to TE junctions defined above were used to generate a binary matrix indicating the presence/absence of the TEs in any given sample. This binary matrix was constructed with custom code based on the observation that there are either many reads or very few reads per sample for any given TE insertion site. After normalizing the number of TE associated reads per sample, a z-score was calculated for every TE across the samples. Positive z-scores were assigned as present and negative z-scores as absent. Because z-score normalization uses the mean of a sample, if all or most of the samples are positive, by definition, half of the samples would end up with a negative z-score. To avoid this mis-identification of positive samples, we add a dummy zero value to the set of samples for every real sample included before z-score calculation. This data was then visualized in R using the gplots function heatmap.2. The identities of blinded samples were estimated based on the clustering of these samples within the dendrogram derived from known samples.

#### Code

Code and notes on running the TE detection and clustering pipeline are available at: https://github.com/mondegreen/DrosCellID.git.

## Results

### Drosophila cells have distinct TE signatures

Previous analysis of available whole genome sequencing (WGS) data revealed that genomic TE distribution can reliably cluster cell lines based on their genotype and laboratory of origin (Han *et al*. 2021). Moreover, WGS analysis using a limited set of six TE families (*297, copia, mdg3, mdg1, roo* and *1731*) was sufficient to replicate the clustering observed when data from all TE families was used (Han *et al*. 2021). Nevertheless, an alternative approach that selectively enriches the six TE families would be more efficient and cost-effective. Therefore, based on these analyses, here we set out to determine if targeted identification of the genomic distribution of a small number of diagnostic TE families could be used to 1) to build an authentication platform for *Drosophila* cell lines based on unique <u>g</u>enomic <u>TE d</u>istribution (gTED) signatures for each cell line, 2) test the validity of this protocol by assessing the identities of blinded samples, both internal and those provided by the *Drosophila* community and 3) assess if cell lines subjected to extensive passaging retain the unique cell-specific gTED signatures.

To achieve these goals, we developed a novel TE based NGS enrichment protocol described in the Materials and Methods (Figure 1). Briefly, this protocol uses a multiplexed nested PCR approach to selectively amplify the library elements containing the 5’ and 3’ ends of the target TE families (Reaction A and B, Figure 1). The products from the final PCR amplification step were subjected to next generation sequencing (NGS) and downstream analyses to determine the type of TE and identify the unique genomic DNA flanking the TE sequence.

The NGS data obtained was first used to identify TE junctions using the bioinformatic strategy outlined in Figure 2. Since the number of reads observed upon amplification with *mdg3*-specific primers was very low, *mdg3* was excluded from further analyses. Normalized counts of reads mapping near TE junctions for the remaining five families were then used to hierarchically cluster all the cell lines. Reads mapping close to the identified TE junctions, whether at 5’ or 3’ end or both, were included in further analyses (Figure 2). The resulting dendrogram showed that the triplicate samples from most cell lines clustering together (Figure 3). Upon processing the NGS data using an alternative approach (Supplementary File 1), a comparable clustering of all the samples was observed (Figure S1). In both approaches, one replicate each from S2 DGRC (S2-DGRC_2) and S2 DRSC (S2-DRSC_2) did not cluster with the other replicates from these cell lines (Figure 3, Figure S1). The similar clustering from both bioinformatic approaches suggests the non-conforming clustering of these two replicates is not an artifact of genomic or computational methods, and was most likely caused by reciprocal sample mislabeling during gDNA extraction. Regardless of the cause of these two discrepancies, the majority of samples (2/3) for both S2 DGRC and S2 DRSC are respectively consistent with one another, providing confidence in the identity of these cell line clusters.

**Figure 3:**
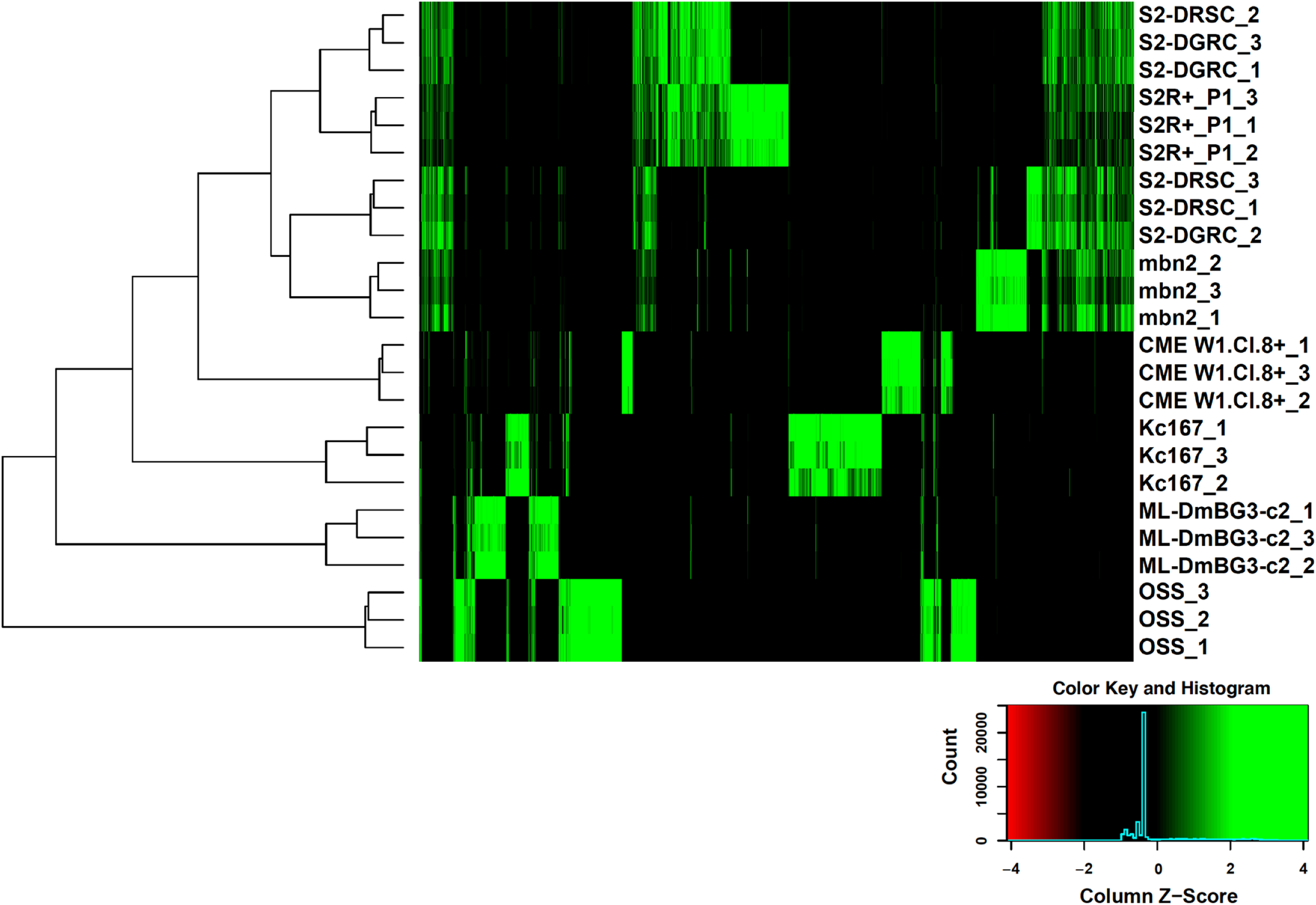
Clustering of cell lines based on genomic transposable element distribution. The cell line clustering was derived upon processing NGS data as described in the Materials and Methods. The triplicates for each cell lines are indicated with 1-3 following the cell line name.

Distinct gTED signatures, a composite of the five TE families assessed, were observed for every cell line investigated (Figure 3 and Figure S2). The tree visualization heatmap demonstrates that there are very few shared TE insertions between all cell lines (Figure 3). In general, the total number of TEs detected by this technique was higher in embryonic cell lines as opposed to cell lines derived from larval or adult tissues (Table 1, Figure S2). The total number of TEs mapped was similar for the replicates of each of the cell lines as seen in the UpSET plot (Lex *et al*. 2014) for these samples (Figure S2). For many of the cell lines, the majority of TE insertions detected were unique relative to those shared with other cell lines. For example, OSS replicates have 262 unique TEs that are not found in any other cell line investigated, with ≤9 TEs in common with any other individual cell lines (Figure S2). The only lines that do not conform to having majority unique TE insertions are S2 DGRC and S2 DRSC as they share a considerable proportion of the TEs with S2R+ (Figure S2). Nevertheless, unique patterns of gTED were sufficient to distinguish between the various S2 sublines (Figures 3, S1 and S2). Two of the three larval tissue derived cell lines (ML-DmBG3-c2, mbn2 and CME W1 Cl.8+) have fewer genomic TE insertions as compared to embryonic S2 and Kc167 lines. However, mbn2, a cell line reportedly derived from the larval circulatory system (Gateff 1977; Gateff *et al*. 1980) has a gTED signature very close to those of the S2 lines, which are all of hematopoietic origin (Schneider 1972). The unexpected similarity between S2 lines and mbn2 was also described recently by Han *et al*. (2021) based on WGS based TE distribution analysis. These analyses demonstrated that the protocol developed to determine genomic distribution of a set of five TE families in *Drosophila* cell lines can be utilized to create unique cell line-specific signatures.

### TE signatures of Drosophila cell lines can be employed for authentication

To assess the value of the developed gTED pipeline and validate it, we next queried if the cell line-specific gTED signatures could be employed to determine the identities of blinded samples (Table 2). The blinded samples were either donations from the *Drosophila* community (external samples) or generated internally at DGRC. All blinded samples, as well as triplicates of an internal control for S2R+ (DGRC_Blinded_control_1-3), were processed as outlined in the Materials and Methods section.

Of the eight external cell lines processed from two different donating labs, six robust gTED signatures were obtained (Figure S3A). However, very few TE insertions detected in six samples, possibly from two cell lines (Figure S3A). gTED profiles for three samples (DRSC_Blinded_13-15) was very similar to the internal control from S2R+ processed in this run (DGRC_Blinded_control_1-3, Figure S3A). For fifteen of the eighteen samples with robust gTED profiles, clusters of triplicates were observed, indicating that each cluster possibly represents replicates samples of five cell lines (Figures 4 and S3A). One sample did not cluster distinctly with any of the other samples (SGLab_Blinded_4, Figures 4 and S3A), however this sample had a gTED profile that is visually most similar to samples SGLab_Blinded_5-6 (Figure S3A). The six samples that had very few TE insertions (triplicates for each labelled DRSC_Blinded_4-6 and DRSC_Blinded_7-9) each passed the genomic DNA and library preparation quality control steps, and the consistent lack of TE insertions among replicates suggested that this was a reproducible signal. Upon clustering the external blinded samples with the previously characterized set of TE signatures it was possible to predict the identities of these samples (Figure 4, Table 2) as DRSC_Blinded_1-3 and DRSC_Blinded_10-12 (Kc167), DRSC_Blinded_4-6 and DRSC_Blinded_7-9 (No identification), DRSC_Blinded_13-15 (S2R+), DRSC_Blinded_16-18 (S2), SGLab_Blinded_1-3 (mbn2) and SGLab_Blinded_5-6 (S2). Moreover, the clustering generated with gTED has the resolution to identify the various S2 sublines. For instance, it is evident that DRSC_Blinded_13-15 are closest to S2R+, DRSC_Blinded_16-18 to S2-DGRC, and SGLab_Blinded_5-6 to S2-DRSC (Figure 4). The investigators who donated the external samples confirmed that the identities determined by the gTED protocol was accurate for all the samples as predicted (Table 2). The two cell lines with very few TE insertions for which a cell line identity prediction could not be generated were mosquito cell lines (Figure 4, Table 2). These experiments demonstrated that the gTED protocol could reliably identify blinded *Drosophila* samples submitted to DGRC by the community.

**Figure 4:**
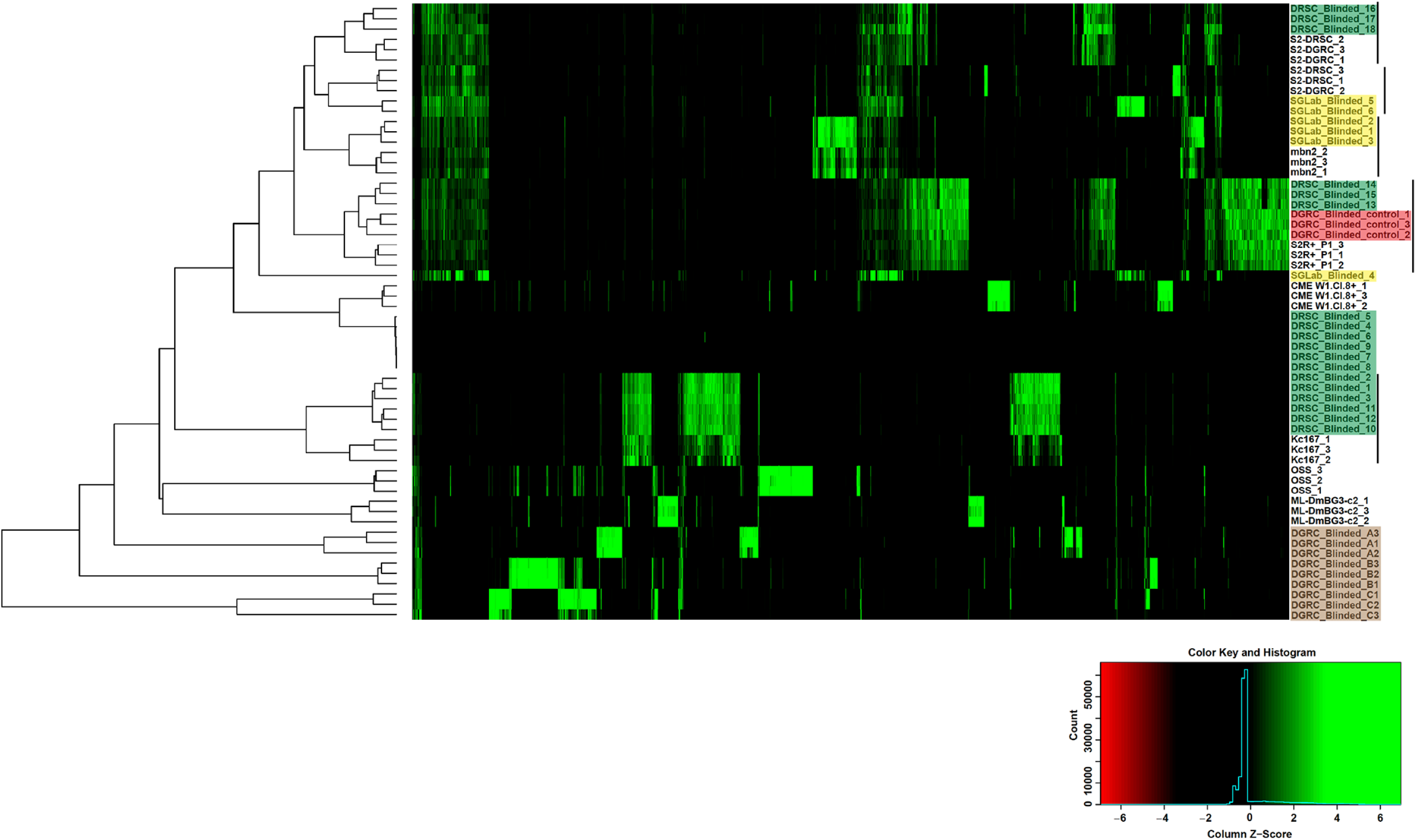
Cell line authentication of double-blind samples using genomic transposable element distribution signatures. Triplicate samples of external blinded cell lines from the lab of Dr. S. Gorski (shaded yellow) and *Drosophila* RNAi Screening Center (shaded green) along with internal blinded samples (shaded brown) and internal control samples (shaded red) were processed with the gTED protocol (Figure 2B) and clustered as described in the Materials and Methods along with the previously processed known samples. The cell lines that the blinded samples cluster with are indicated with the black lines. Internal blinded samples cluster as a separate group. Samples DRSC_Blinded_4-9 with very few or no TEs detected were from mosquito cell lines (Table 2).

All three internal blinded cell lines had unique gTED signatures that clustered distinctly relative to all previously-characterized gTED signatures (Figures 4 and S3B). Nevertheless, the triplicates from each of the internal blinded cell lines reliably clustered together (Figure 4). Upon unblinding (Table 2), the internal blinded samples were found to be from three cell lines not included in the initial development phase of the project: 1182-4H (DGRC_Blinded_A, DGRC#177, CVCL_Z708), Ras[V12];wts[RNAi] (DGRC_Blinded_B, DGRC#189, CVCL_IY71) and delta_l(3)mbt-OSC (DGRC_Blinded_C, DGRC#289). Thus, processing blinded samples through the gTED pipeline revealed that 1) reliable identification of samples with known gTED signatures can be achieved, 2) the protocol is capable of distinguishing *Drosophila* versus non-*Drosophila* cell lines and 3) *D. melanogaster* cell lines previously uncharacterized by the gTED protocol can be identified as such, without providing a false identification.

### TE signature of S2R+ is retained upon extensive passaging

Extensive passaging of cell lines can potentially alter cellular genomes (Wenger *et al*. 2004; Oh *et al*. 2013). Apart from gross genomic changes, extensive passaging introduced single nucleotide polymorphisms in mammalian cell lines (Pavlova *et al*. 2015). To determine the effect of extensive passaging on the gTED signatures generated in this study, we passaged S2R+ cell line 50 times and isolated genomic DNA in triplicate at every tenth passage for processing (Fig. 5A). Upon generating a cluster using the gTED protocol, it is evident that the triplicates from the passages cluster randomly and not according to passage numbers (Fig. 5B). Moreover, all replicates from every passage tested form a distinct cluster (Fig. S4) indicating that extensive passaging of S2R+ does not alter the S2R+ gTED signature for up to 50 passages.

**Figure 5:**
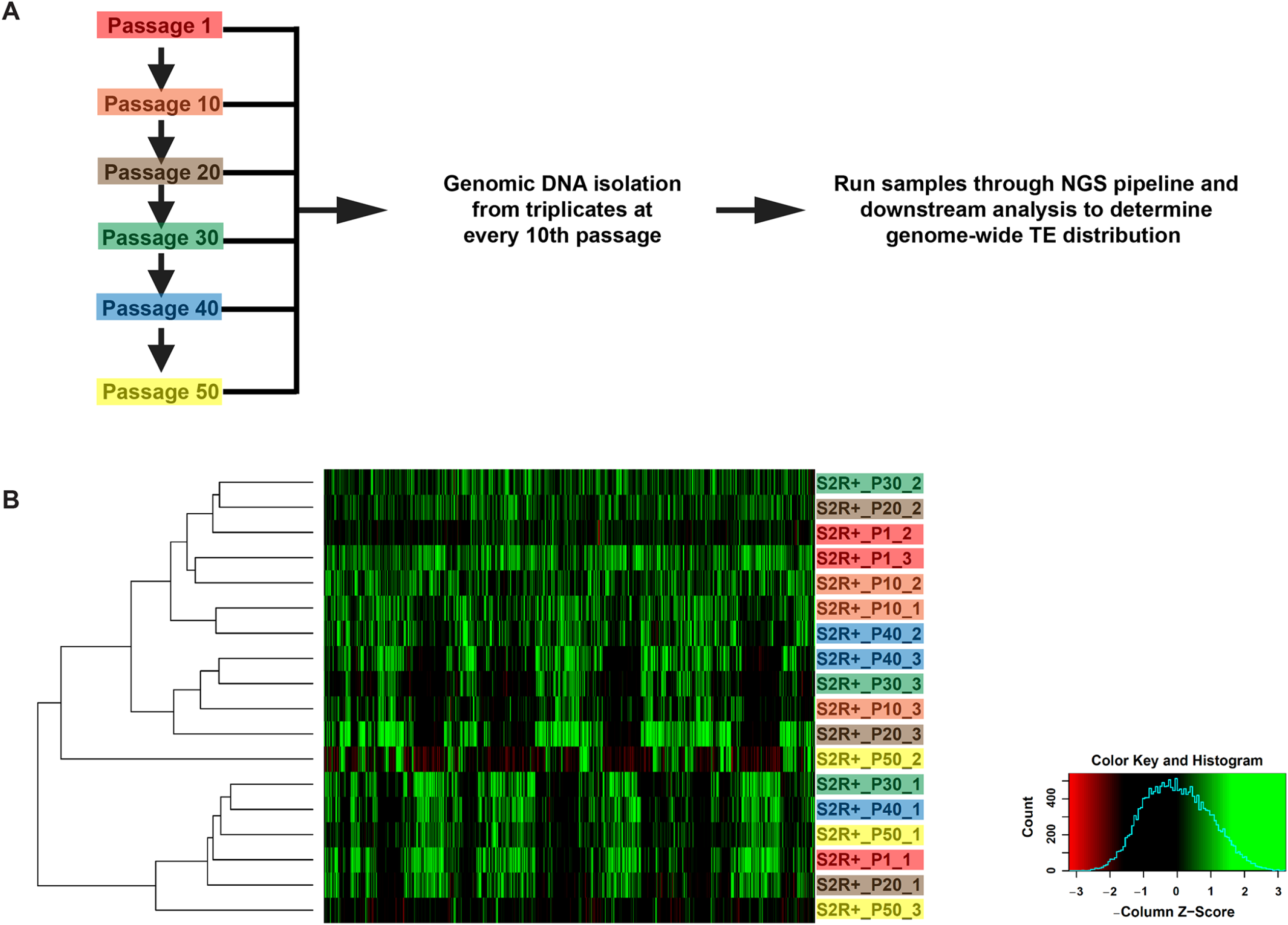
Genomic transposable element distribution signatures for S2R+ cells do not cluster by passage number. **A**) Schematic outlining the protocol to acquire samples between 1-50 S2R+ passages for assessment by the gTED protocol. **B**) Clustering of all the passage samples generated based on TE predictions. The triplicates samples of every passage are shaded in one color each.

## Discussion

The aim of this study was to develop and test a cell authentication protocol that could reliably identify the most commonly used *Drosophila* cell lines to help researchers validate their reagents as per the NIH mandate. Our novel protocol allowed us to define unique gTED signatures that could identify each of the *Drosophila* cell lines that were tested here. In addition, the resolution obtained from the gTED signatures allows for distinguishing between S2 sublines. Data presented here demonstrate that the gTED signatures of the replicates of most cell lines cluster together, outlining the reproducibility of the gTED protocol while also underscoring the value of having replicate samples for reliable cell line identification. Crucially, accurate identification of blinded samples donated by the research community validated the gTED protocol in a real-world setting.

To reliably identify a *D. melanogaster* cell line using the gTED protocol, an established gTED signature is a prerequisite. Towards this end, we have now established gTED signatures for the widely distributed lines, S2R+, S2 DGRC, S2 DRSC, Kc167 and ML-DmBG3-c2 lines (Luhur *et al*. 2018). In addition, gTED signatures are also available for OSS, mbn2, CME W1 Cl.8+, 1182-4H, Ras[V12];wts[RNAi] and delta l(3)mbt-OSC lines. Importantly, the lack of an established gTED signature does not lead to misidentification, as was observed with the internal blinded samples. In the event that a cell line without an established gTED signature needs to be authenticated, a stock from the DGRC repository with the same identity will be assayed concurrently to serve as a control. In due course, DGRC will also expand the gTED protocol to include as many cell lines from our repository as possible. These efforts will ensure the creation of a comprehensive database of gTED signatures for *Drosophila* cell lines.

Mosquito cell lines included as blinded samples helped clarify that the gTED protocol can discriminate non-*Drosophila* cell lines. In *Ae. aegypti* and *An. gambiae*, 10% and 6% of the total genome, respectively, is comprised of LTR retrotransposons (Nene *et al*. 2007; Melo and Wallau 2020). Presence of active LTR transposons, specifically *Ty1/copia* has also been described in Aag2 (*Ae. aegypti*) cells (Maringer *et al*. 2017). Since we confirmed that the DNA and library preparation for these samples were comparable, it is most likely therefore that the TE-specific primers used in this study cannot amplify mosquito TE families. Our results demonstrate that in pure samples mosquito cells can be distinguished from *D. melanogaster* cell lines using the gTED protocol. However, detecting low levels of inter- or intra-species contamination might a more challenging pursuit. A *D. melanogaster* cell line contaminated with low levels of a mosquito cell line is unlikely to be detected with gTED, necessitating using other methods for such specific instances. A future avenue is to explore the sensitivity of the gTED protocol to intra- or inter-species contamination. In addition, it will be imperative to determine if we can determine low levels of contamination of *Drosophila* cell lines containing unique gTED signatures.

Our analysis also demonstrated that the genomic distribution of TEs is largely unchanged over 50 passages in S2R+ cells. The narrow window into the passaging-associated genomic structure provided by the gTED protocol is most likely not representative of more complex genomic and/or transcriptomic changes that the extensively passaged cells might have undergone. Nevertheless, S2R+ cells passaged continuously for up to 50 times can still be identified with the gTED protocol. Among the S2 lines assessed in this study, it has been proposed that the S2R+ line is possibly the closest to the original Schneider line (Schneider 1972; Yanagawa *et al*. 1998). The other two S2 sublines, S2-DGRC and S2-DRSC, are isolates with less clear history from the original Schneider isolates before being added to the DGRC repository (Ayer and Benyajati 1992; Cherry *et al*. 2005). All three of the S2 sublines assessed have unique gTED signatures that discriminate them and can be used to identify blinded cell lines precisely to the S2 subline. In general, S2 sublines have a more complex TE-landscape, higher aneuploidy and copy number variation than other *D. melanogaster* cell lines (Han *et al*. 2021). The possibility that the gTED signature can be used as a proxy for broader genomic changes remains to be investigated.

In summary, utilizing the genomic distribution of five TE families we have developed the gTED pipeline to facilitate the authentication of *Drosophila* cell lines. We demonstrate that the developed gTED protocol can assign distinct signatures to the various *Drosophila* cell lines tested. Blinded and extensively passaged samples can now be authenticated employing the gTED protocol. Researchers working with *Drosophila* cell lines can independently authenticate cell lines being used in their laboratories using the protocol and code described in this study. Alternatively, DGRC will implement a cost-based service for the research community to access and authenticate their cell lines for both publications and research funding. Ultimately, our goal is to include more cell lines from the DGRC repository into the gTED pipeline and generate gTED signatures for all cell lines deposited with the DGRC.

## Data availability

All data necessary for confirming the conclusions in this paper are included in this article and in supplemental figures and tables. All the NGS data has been deposited at Sequence Read Archive available with the accession number: SRP323476

## Acknowledgments

We thank Dr. Kris Klueg for critical input in shaping the manuscript. We thank Grace Kim (DRSC), Dr. Stephanie Mohr (DRSC), Nancy Erro Go (British Columbia Cancer Research Centre) and Dr. Sharon Gorski (British Columbia Cancer Research Centre) for providing the blinded genomic DNA samples. We thank Chunlin Yang, Jie Huang, and Sumitha Nallu at the Center for Genomics and Bioinformatics (Indiana University, Bloomington) for their help with NGS sample processing. This work was supported by NIH grants (5P40OD010949 and 3P40OD010949-16S2) to the *Drosophila* Genomics Resource Center, National Science Foundation under Grant No. CNS-0521433 to Center for Genomics and Bioinformatics (Indiana University, Bloomington) and the University of Georgia Research Foundation to CMB.

## Figure Legends

***Supplementary Figure 1***: **Clustering of cell lines based on genomic transposable element distribution using an alternative bioinformatics pipeline**. The cell line clustering is derived from processing NGS data as described in Supplementary File 1. The triplicates for each cell lines are indicated with 1-3 following the cell line name.

***Supplementary Figure 2***: **Unique TEs distinguish cell lines assessed by gTED**. The number of TEs that are shared between the samples (Intersection size) are plotted in this UpSET plot. Filled in dots indicate the samples that share the particular set of TEs. The absolute number of TEs for each of the samples is plotted as Set Size.

***Supplementary Figure 3***: **Blinded samples have unique gTED signatures**. External (**A**) and internal (**B**) blinded samples assessed using the gTED protocol have unique gTED signatures that cluster replicates by cell identity.

***Supplementary Figure 4***: **S2R+ cells retain unique gTED signature despite extensive passaging**. All samples from this study assessed using the gTED protocol indicate that all the S2R+ passages cluster together, still retaining a unique cell-line specific gTED signature. The S2R+ passages are shaded in green.

***Supplementary File 1***: **Description of the alternative bioinformatics pipeline used to cluster cell lines based on genomic transposable element distribution**. Clustering using this alternative approach for cell lines used in the development phase of the project is shown in Supplementary Figure 1.

***Supplementary File 2***: **Table of samples ID listed in SRA accession used for gTED analysis**. The 75 samples used for the analysis in the manuscript are listed in the table. The other 39 samples listed in SRP323476 were used for testing and development.

***Supplementary File 3***: **Presence absence matrix for cell line clustering**. The final data matrix used for cell line clustering is available at: https://github.com/mondegreen/DrosCellID/blob/main/combined.presence-absence.example.tsv.

